# Type I interferon signaling enhances kainic acid-induced seizure severity

**DOI:** 10.1101/2024.11.13.623521

**Authors:** Jeong-Hwa Ma, Jun-Cheol Eo, Changjun Lee, Inhwa Hwang, Jihye Choi, Sung Jae Shin, Chul Hoon Kim, Je-Wook Yu

## Abstract

Epilepsy is a chronic neurological disorder characterized by recurrent seizures, yet the role and mechanisms of type I interferon (IFN) signaling in seizure conditions remain elusive. In this study, we demonstrate that type I IFN signaling exacerbates seizure phenotypes in a kainic acid-induced seizure mouse model. We found that the absence of type I IFN signaling in *Ifnar1*^-/-^ mice led to decreased neuronal excitability and microglial activation in response to kainic acid stimulation. Conversely, intracerebroventricular injection of IFN-β heightened the severity of kainic acid-induced seizures. *In vitro* calcium imaging revealed that IFN-β treatment amplified both basal and kainic acid-induced neuronal excitability, though no significant difference was observed in basal neuronal excitability between wild-type and *Ifnar1*^-/-^ neurons. Furthermore, *Ifnar1*^-/-^ mice exhibited reduced mTOR activation in the brain following kainic acid administration. Consistent with this finding, IFN-β treatment induced mTOR activation, as indicated by S6 phosphorylation in *in vitro* mixed glial cultures. Taken together, these results demonstrate the critical role of type I IFN signaling in seizure pathogenesis and suggest that targeting type I IFNs could be a promising therapeutic strategy for reducing seizure severity and mitigating epilepsy.

## Introduction

Epilepsy is a chronic neurological disorder characterized by recurrent seizures, affecting over 65 million people worldwide^1,2^. It can result from genetic mutations impacting ion channel function (*SCN1A*, *KCNQ2/3*), synaptic transmission (*LGI1*, *STXBP1*), or metabolic pathways (*SLC2A1*, *TSC1/2*), as well as from interactions between genetic predisposition and environmental factors^3–5^. These genetic and environmental contributors collectively lead to neuronal hyperexcitability, resulting in abnormal electrical activity in the brain that manifests as seizures and ultimately develops into epilepsy^6^. Despite advancements in anti-epileptic drugs designed to target neuronal excitability, nearly one-third of patients remain refractory to these treatments^7–9^. Moreover, the mechanisms underlying epileptogenesis**–**the gradual process by which a normal brain becomes epileptic**–**are still not well understood^10^. These challenges highlight critical gaps in our understanding of epilepsy and underscore the need for deeper investigation into the pathological processes and molecular changes involved in epileptogenesis.

Type I interferon (IFN) signaling has emerged as a key factor in the development and maintenance of the central nervous system (CNS), playing crucial roles in neural development, astrocyte-synapse regulation, and microglial phagocytosis^11–15^. In the CNS, various cell types, including microglia, produce type I IFN, and its receptors, IFNAR1 and 2, are widely expressed across most brain cell types^11,15,16^. Importantly, dysregulation of type I IFN signaling has been associated with multiple CNS diseases, including Alzheimer’s disease, Down syndrome, and Parkinson’s disease, highlighting its crucial role in CNS function^17–26^.

Although type I IFN signaling has been studied in the context of various CNS diseases, its role in seizure conditions remains largely unexplored. A recent study has shown that type I IFN signaling is upregulated in the microglia of seizure-induced mice^27^. Additionally, clinical reports indicate that spontaneous seizures frequently occur during interferon therapy^28,29^. Moreover, studies on poly (I:C)-mediated TLR3 activation and IFN treatment have produced conflicting results regarding its influence on neuronal excitability, with evidence of both beneficial and detrimental effects^12,30–32^. Collectively, these findings suggest that type I IFNs may play a role in the initiation and progression of seizures. However, the specific involvement and mechanisms of type I IFN signaling in seizure conditions remain unclear. Therefore, in the present study, we aimed to investigate the role of type I IFN signaling under seizure conditions.

## Materials and methods

### Mice

C57BL/6 mice were obtained from Orient Bio. *Ifnar1*^-/-^ mice (JAX 32045) were kindly gifted from Dr. Heung Kyu Lee (KAIST, Republic of Korea). For primary neuron culture, pregnant mice were purchased from Koatech. All mice were maintained under specific pathogen-free conditions at Yonsei University College of Medicine. 8- to 12-week-old male mice were used for the experiments. All animal experimental protocols were approved by the Institutional Ethical Committee, Yonsei University College of Medicine (2022-0121). All experiments were conducted in accordance with the approved guidelines of the Institutional Ethical Committee.

### Mice treatment

To induce seizures, mice received intraperitoneal injections of kainic acid (20-21 mg/kg) dissolved in saline^33,34^. Seizure behavior was monitored for 2 h at 10 min intervals using the modified Racine scale^35^: (0) no response; (1) freezing behavior; (2) head nodding; (3) forelimb clonus; (4) rearing and falling; (5) loss of posture and generalized convulsive activity; (6) death. For intracerebroventricular administration, mice were anesthetized with an intraperitoneal injection of Zoletil (tiletamine/zolazepam, 40 mg/kg) and Rompun (xylazine, 10 mg/kg), and then placed on a stereotaxic apparatus. A 26-gauge stainless steel cannula (Plastic one Inc.) was inserted using the following coordinates to target the right lateral ventricle: AP -0.5 mm, ML +1.0 mm, and DV -2.0 mm. After surgery, mice were allowed to recover for at least one week.

Following the recovery period, recombinant IFN-β (50 μg/ml) or vehicle (1 mg/ml BSA in PBS) was administered at a volume of 1 μl at a rate of 1 μl/min for 3 consecutive days. Six hours after the final injection, kainic acid was administered. For immunoblotting experiments, mice were anesthetized, transcardially perfused with PBS, and brain tissues were isolated after removing olfactory regions, brainstem, and meninges.

### Reagents and antibodies

Kainic acid (7065) and rapamycin (1292) were purchased from Tocris Bioscience. Recombinant mouse IFN-β was obtained from R&D Systems (8234-MB) and Fluo-3AM was purchased from Invitrogen (F23915). For immunohistochemistry, anti-c-FOS (Cell Signaling Technology, 2250), anti-IBA1 (Wako, 019-19741), and anti-GFAP (Invitrogen, 13-0300) antibodies were used. For immunoblotting, anti-phospho-S6 (4858), anti-S6 (2217), anti-pAKT (9271), anti-AKT (9272), anti-phospho-STAT1 (9167), and anti-STAT1 (14994) antibodies were purchased from Cell Signaling Technology. Anti-β-actin antibody was obtained from Santa Cruz Biotechnology (sc-47778). For flow cytometry, anti-CD45-BV421 (BD, 563890), anti-CD11b-PE (Invitrogen, 12-0112-82), and anti-ACSA2-APC (Miltenyi, 130-117-535) antibodies were used.

### Immunohistochemistry

Mice were deeply anesthetized with isoflurane (5% in O_2_) and transcardially perfused with PBS, followed by 4% paraformaldehyde (PFA). The dissected brains were post-fixed in 4% PFA and subsequently cryoprotected in 30% sucrose at 4 °C. The brains were sectioned into 40 μm slices using a Leica CM1860 cryostat. The sections were blocked and permeabilized in a blocking solution containing 4% BSA and 0.3% Triton X-100 at room temperature for 60 min. They were then incubated with primary antibodies diluted in the blocking solution for 24– 48 h at 4 °C. After several washes, the sections were incubated with Cy3- or Alexa Fluor 488- conjugated secondary antibodies in the dark. Following additional washes, nuclei were stained with DAPI. Z-stack images were acquired at 0.75 μm intervals over a 30 μm Z-range using confocal microscopy (LSM980, Carl Zeiss). The Z-stack images were processed into single-plane maximal intensity projections using Zen Blue software (Carl Zeiss).

For microglia morphology analysis, microglia in Z-projection images were classified into three categories based on the ratio of soma diameter to the length of the longest process: Ramified (process length greater than 3 times the soma diameter), Bushy (process length between 1.5 and 3 times the soma diameter), and Amoeboid (process length less than 1.5 times the soma diameter). A total of 10–15 microglia were randomly selected from each sample, and the percentage of microglia in each category was calculated. For mean fluorescence intensity (MFI) analysis, ImageJ (NIH) software was used. Fluorescence intensity was normalized to DAPI staining to determine the MFI.

### Primary neuronal culture and calcium imaging

Cortical neurons were prepared from embryonic day 17-18 mice. Cortices were dissected in a dissection solution containing 1% HEPES (Gibco, 15630080) and 1% Penicillin-Streptomycin (Gibco, 15070063) in HBSS (Gibco, 14175095). The tissue was incubated in 0.05% trypsin (Gibco, 15090046) at 37 °C for 12 min. Following trypsinization, cortices were gently triturated in a chopping solution containing 600 U/ml Deoxyribonuclease Ⅰ (Sigma, D5025) in dissection solution to dissociate the cells. The cells were then washed with Neurobasal medium (Gibco, 21103049) and plated onto poly-D-lysine-coated confocal dishes. Neurons were cultured in Neurobasal medium supplemented with 2% B-27 (Gibco, 17504001) and 1% Glutamax (Gibco, 35050061) for 14 days, with medium replaced every 2–3 days.

For calcium imaging, primary neurons were incubated with 3 μM Fluo-3 AM calcium indicator for 30 min at 37 °C. Then, live calcium imaging was performed using an LSM980 confocal microscope, acquiring time-lapse images at a rate of 0.6 frames per second over 8 min. The fluorescence intensity (MFI) of Fluo-3 AM was normalized to the baseline fluorescence (F0), calculated from the first 20 frames. Following baseline acquisition, neurons were treated with 1 mM KA and 50 mM KCl to assess neuronal excitability in response to those stimuli. Only neurons that exhibited a calcium response to KCl were included in the analysis. For the IFN-β priming experiment, initial baseline fluorescence was recorded for 2 min, followed by the addition of 5 ng/ml recombinant IFN-β to the media. Neurons were incubated with IFN-β for 30 min at 37 °C. After incubation, neuronal excitability was measured for 0.5 min, after which KA and KCl were applied. The effect of IFN-β priming was evaluated by comparing the neuronal excitability of IFN-β-primed neurons to that of unprimed controls.

### Mixed glia culture

Mixed glial cells were isolated from the whole brains of postnatal day 0-2 mice as previously described^36^. Briefly, the cerebellum and olfactory bulbs were removed, and the meninges were carefully peeled off in Hank’s balanced salt solution. The brain tissue was then homogenized by gentle pipetting using Pasteur pipettes of progressively smaller diameters. The resulting homogenate was centrifuged and passed through 70 μm cell strainer to remove debris. The dissociated cells were plated onto T75 flaks in DMEM/F12 (1:1) medium supplemented with 10% FBS and antibiotics. Cells were cultured for 14 days, with the medium changed after 7 days and subsequently every 2 days. The mixed glial population was analyzed by flow cytometry (FACSVerse, BD) using FlowJo software (TreeStar).

### Immunoblotting

Total proteins were extracted from the hippocampus or cortex using RIPA buffer containing 50 mM Tris-Cl (pH 8), 150 mM NaCl, 0.1% SDS, 1 mM EDTA, 1% Triton X-100, 0.05% sodium deoxycholate, and protease inhibitors. Mixed glial cells were lysed in buffer containing 20 mM Tris-Cl (pH 7.5), 0.5% Nonidet P-40, 50 mM KCl, 150 mM NaCl, 1.5 mM MgCl_2_, 1 mM EGTA, and protease inhibitors. Protein concentrations were determined using the Bradford assay. Samples were separated by 10% SDS-polyacrylamide gel electrophoresis and transferred to PVDF membranes. The membranes were briefly washed with PBST and blocked with 3% skim milk at room temperature for 1 h, followed by incubation with primary antibodies. After washing with PBST, the membranes were incubated with horseradish peroxidase-conjugated secondary antibodies for 1–2 h at room temperature. Following additional washes, the antibodies were visualized using Pierce™ ECL Western Blotting Substrate (Thermo Fisher Scientific, 32209). Blot visualization was performed using the ImageQuant™ 800 system, and images were cropped for presentation. All blots were replicated in at least three independent experiments. Protein levels were quantified using ImageJ software.

### Statistical analysis

All data are presented as the mean and standard error of the mean (SEM) of individual samples. The Mann-Whitney *U* test was used for seizure score comparisons between wild-type and *Ifnar1^-/-^* mice. Other analyses were conducted with Student’s *t* test or two-way analysis of variance (ANOVA), followed by Tukey’s post hoc test to explore interactions between the factors. P-values < 0.05 were considered statistically significant. All statistical analyses were performed using Prism 9.0 (GraphPad Software).

## Results

### Deficiency of type Ⅰ interferon signaling reduces seizure severity in kainic acid-induced seizures

Through analysis of publicly available RNA-seq data, we observed an upregulation of several interferon-stimulated genes in microglia from a kainic acid (KA)-induced seizure mouse model (Supplementary Fig. 1A, B) and in the hippocampus of pilocarpine-induced seizure mouse model (Supplementary Fig. 1C, D)^27,37^. To evaluate the role of type I IFN signaling in seizure phenotypes, we induced seizures in wild-type and *Ifnar1^-/-^* mice using KA and compared seizure severity between the two groups (Fig. 1A). Notably, *Ifnar1^-/-^* mice exhibited significantly reduced seizure severity compared to wild-type mice (Fig. 1B). We also monitored body weight for 7 days post-seizure as an indicator of recovery from seizure-induced aftereffects. Wild-type mice showed more severe and prolonged weight loss than *Ifnar1^-/-^* mice (Fig. 1C), indicating that the absence of type I IFN signaling mitigates the long-term impact of seizures.

**Fig 1.**
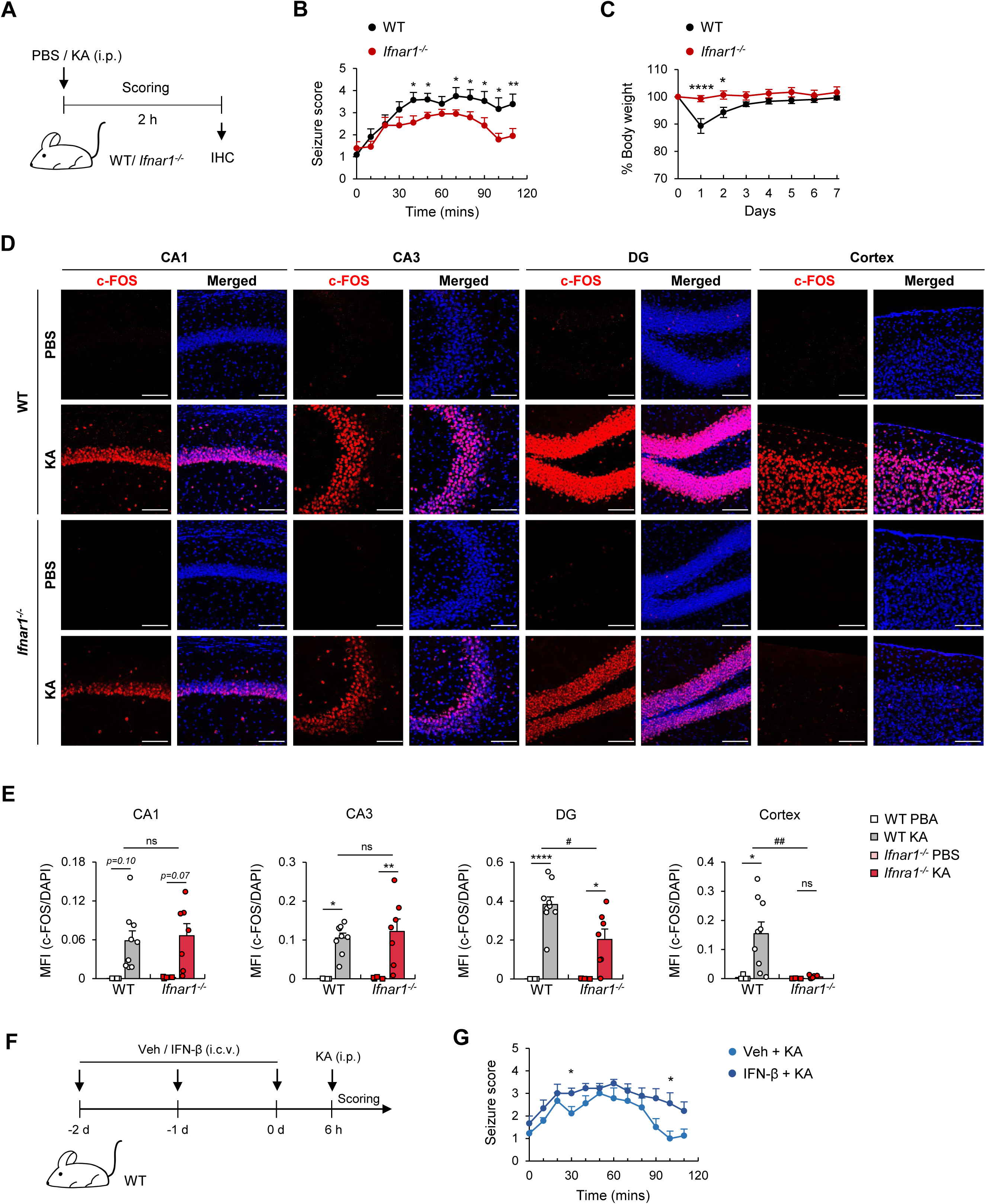
Kainic acid injection induces generalized seizures with widespread neuronal activation in wild-type (WT) mice, but focal seizures with limited neuronal activation in *Ifnar1^-/-^* mice. **(A)** Schematic diagram for kainic acid (KA)-induced seizure. Mice were injected with either PBS or KA (21 mg/kg, i.p.), monitored for seizure activity using a modified Racine scale for 2 h, and harvested for immunohistochemistry. **(B)** Seizure scores in WT and *Ifnar1^-/-^* mice post-KA injection. WT, *n* = 21; *Ifnar1^-/-^*, *n* = 23. **(C)** Body weight changes in WT and *Ifnar1^-/-^* mice after KA administration. WT, *n* = 6; *Ifnar1^-/-^*, *n* = 5. **(D)** Representative images of c-FOS (red) and DAPI (blue) staining in the hippocampal subregions CA1, CA3, and dentate gyrus (DG), as well as the cortex, of PBS- or KA- injected WT and *Ifnar1^-/-^* mice. Scale bar, 100 μm. **(E)** Quantification of mean fluorescence intensity (MFI) of c-FOS+ signals normalized to DAPI across brain regions. PBS, *n* = 4; KA, *n* = 7-9. **(F)** Schematic diagram for KA-induced seizure following IFN signaling priming. Mice were injected with either vehicle or IFN-β (50 ng, i.c.v.) once daily for 3 days. 6 h after the final injection, KA (20 mg/kg, i.p.) was administered, and seizure activity was measured for 2 h. **(G)** Seizure scores in vehicle- and IFN-β-injected mice after KA-injection. *n* = 9 per group. Data are presented as mean ± SEM. Statistical analysis was performed using the Mann-Whitney *U* test for seizure scores (B, G), Student’s *t* test for body weight changes (C), and two-way ANOVA with Tukey’s post-hoc test for c-FOS expression (E). *,^#^ *P* < 0.05, **,^##^ *P* < 0.01, **** *P* < 0.0001, ns; not significant.

To investigate whether the observed behavioral differences were linked to variations in neuronal activation, we assessed c-FOS expression, a marker of neuronal activation, in the brains of wild-type and *Ifnar1^-/-^* mice 2 h after KA administration. PBS-injected controls exhibited minimal c-FOS expression, while KA administration led to elevated c-FOS levels in both wild-type and *Ifnar1^-/-^* mice. Although no significant differences were observed in the mean fluorescence intensity (MFI) of c-FOS in the CA1 and CA3 regions of the hippocampus, c-FOS expression was significantly lower in the dentate gyrus (DG) and cortex of *Ifnar1^-/-^* mice compared to wild-type mice (Fig. 1D, E). Given that KA predominantly induces focal seizures in the temporal lobe^33^, these results suggest that neuronal activation in wild-type mice extended beyond the hippocampus to the cortex, whereas in *Ifnar1^-/-^* mice, activation remained largely confined to the hippocampus.

To confirm that the difference in seizure severity was specifically attributable to the absence of type I IFN signaling, we administered IFN-β, a type I interferon, into the lateral ventricles of wild-type mice for 3 consecutive days to enhance interferon signaling. Six hours after the final injection, KA was administered, and seizure behavior was subsequently monitored (Fig. 1F). Consistently, IFN-β-treated mice exhibited slightly more severe seizures than vehicle-treated controls, further confirming that type I IFN signaling levels can modulate neuronal activity and seizure severity (Fig. 1G).

### *Ifnar1^-/-^* mice exhibit distinct microglial morphology during kainic acid-induced seizures

Microglia are known to undergo phenotypic changes in response to epileptic stimuli^38–42^. To investigate whether the reduced neuronal activity observed in *Ifnar1^-/-^* mice is associated with altered microglial activity, we compared the mean fluorescence intensity (MFI) of IBA1, a microglial marker, and microglial morphology in brain sections from wild-type and *Ifnar1^-/-^* mice 2 h after KA administration (Fig. 2A). Although both wild-type and *Ifnar1^-/-^* mice exhibited an increasing tendency in IBA1 MFI following KA treatment, no significant differences were observed between the genotypes in hippocampal regions CA1 and CA3. However, in the cortex, wild-type mice showed a marked increase in IBA1 MFI in response to KA, whereas *Ifnar1^-/-^* mice did not display a significant change, suggesting a correlation between neuronal activation and the microglial response during KA-induced seizures (Fig. 2B, C).

**Fig 2.**
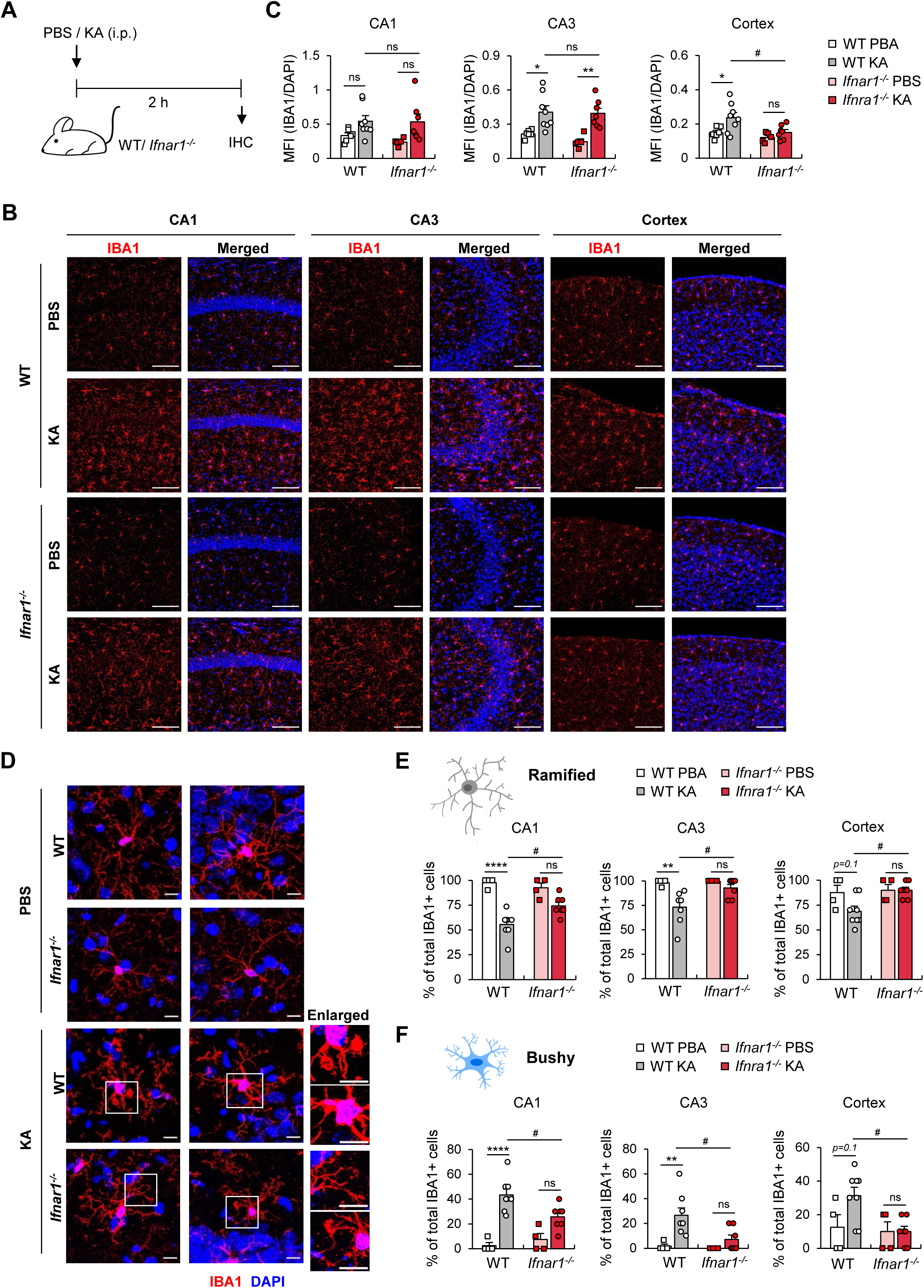
Kainic acid-induced seizures elicit distinct microglial responses in WT and *Ifnar1^-/-^* mice. **(A)** Schematic of kainic acid-induced seizures. **(B)** Representative images of IBA1 (red) and DAPI (blue) staining in the hippocampal subregions CA1, CA3, and cortex, from PBS- or KA- injected WT and *Ifnar1^-/-^* mice. Scale bar, 100 μm. **(C)** Quantification of MFI of IBA1+ signals normalized to DAPI across brain regions. PBS, *n* = 5-6; KA, *n* = 7-8. **(D)** Magnified images showing detailed morphology of IBA1+ microglia. Scale bar, 10 μm. **(E, F)** Percentage of microglia displaying ramified or bushy morphology across treatment groups. Data are presented as mean ± SEM. Statistical analysis was performed using two-way ANOVA with Tukey’s post-hoc test for (C, E, F). *,^#^ *P* < 0.05, ** *P* < 0.01, **** *P* < 0.0001, ns; not significant.

We next examined changes in microglial morphology under seizure conditions. Following KA stimulation, microglia in the wild-type mice exhibited phagocytic bodies and thickened processes, whereas these phenotypes were rarely observed in *Ifnar1^-/-^* mice (Fig. 2D). To further evaluate microglial activation, we categorized microglia into three morphological subsets: ramified, bushy, and amoeboid (Fig. 2E, F and Supplementary Fig. 2). In wild-type mice, significant morphological changes were observed, with microglia transitioning from a ramified to a bushy form upon KA stimulation, whereas these changes were minimal in *Ifnar1^-/-^* mice in both the hippocampus and cortex (Fig. 2E, F). Amoeboid microglia were rarely detected in either genotype (Supplementary Fig. 2). In contrast to the changes in microglial intensity and morphology, the intensity of GFAP, an astrocyte marker, was not significantly affected by KA treatment in either wild-type or *Ifnar1^-/-^* mice (Fig. 3A, B). These findings suggest that type I IFN signaling plays a critical role in modulating microglial activation and morphology during seizure conditions.

**Fig 3.**
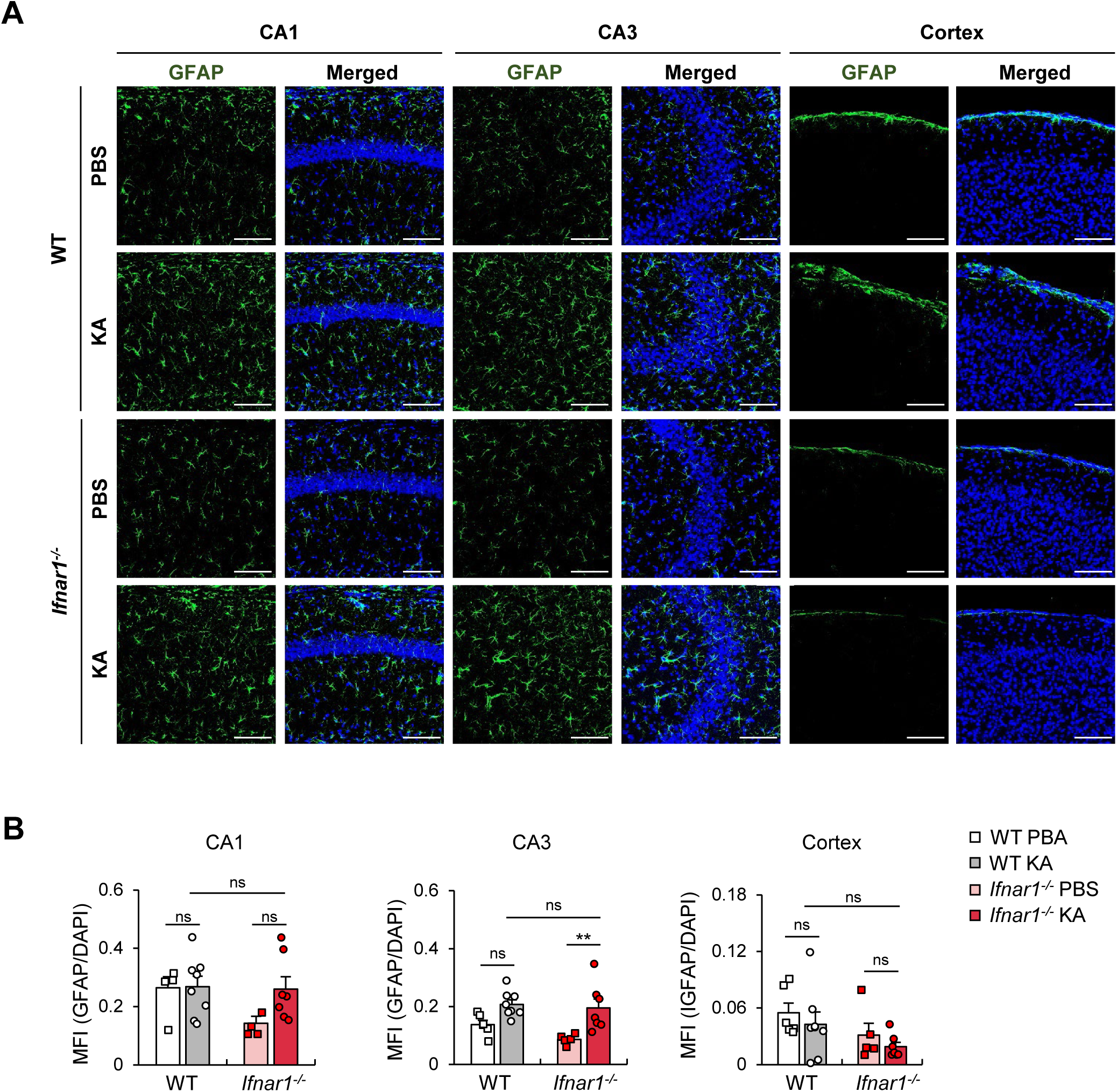
Kainic acid-induced seizures do not significantly impact astrocyte activation in WT and *Ifnar1^-/-^* mice. **(A)** Representative images of GFAP (green) and DAPI (blue) staining in the hippocampal subregions CA1, CA3, and cortex, from PBS- or KA- injected WT and *Ifnar1^-/-^* mice. Scale bar, 100 μm. **(B)** Quantification of MFI of GFAP+ signals normalized to DAPI across brain regions. PBS, *n* = 5-6; KA, *n* = 7-8. Data are presented as mean ± SEM. Statistical analysis was performed using two-way ANOVA with Tukey’s post-hoc test. ** *P* < 0.01, ns; not significant.

### Type I interferon can directly influence neuronal activation

We next investigated whether type I IFN can directly modulate neuronal activity by performing calcium imaging on primary neurons to measure their response to stimuli. Neurons were loaded with the calcium indicator Fluo-3 AM to detect calcium influx, a hallmark of neuronal excitability. Changes in neuronal excitability were quantified as the ratio of Fluo-3 AM MFI to baseline MFI (ΔF/F_0_). Baseline activity (F_0_) was recorded before neurons were sequentially treated with KA to induce excitability, followed by KCl to confirm cell responsiveness (Fig. 4A). Both KA and KCl treatments successfully triggered calcium influx, as demonstrated by increased fluorescence signals (Fig. 4B). Since KCl-induced fluorescence signals were substantially larger than those induced by KA, overshadowing the KA response in subsequent analyses, KCl treatment was omitted from the experimental schematic and data presentations, though it was consistently performed. Consequently, only KA-induced fluorescence changes are emphasized in the following analyses. We first compared KA-induced neuronal responses between wild-type and *Ifnar1^-/-^* neurons. The maximum calcium influx amplitude, defined as the peak ΔF/F_0_ induced by KA, was assessed, and no significant differences were observed between the two genotypes (Fig. 4C–E).

**Fig 4.**
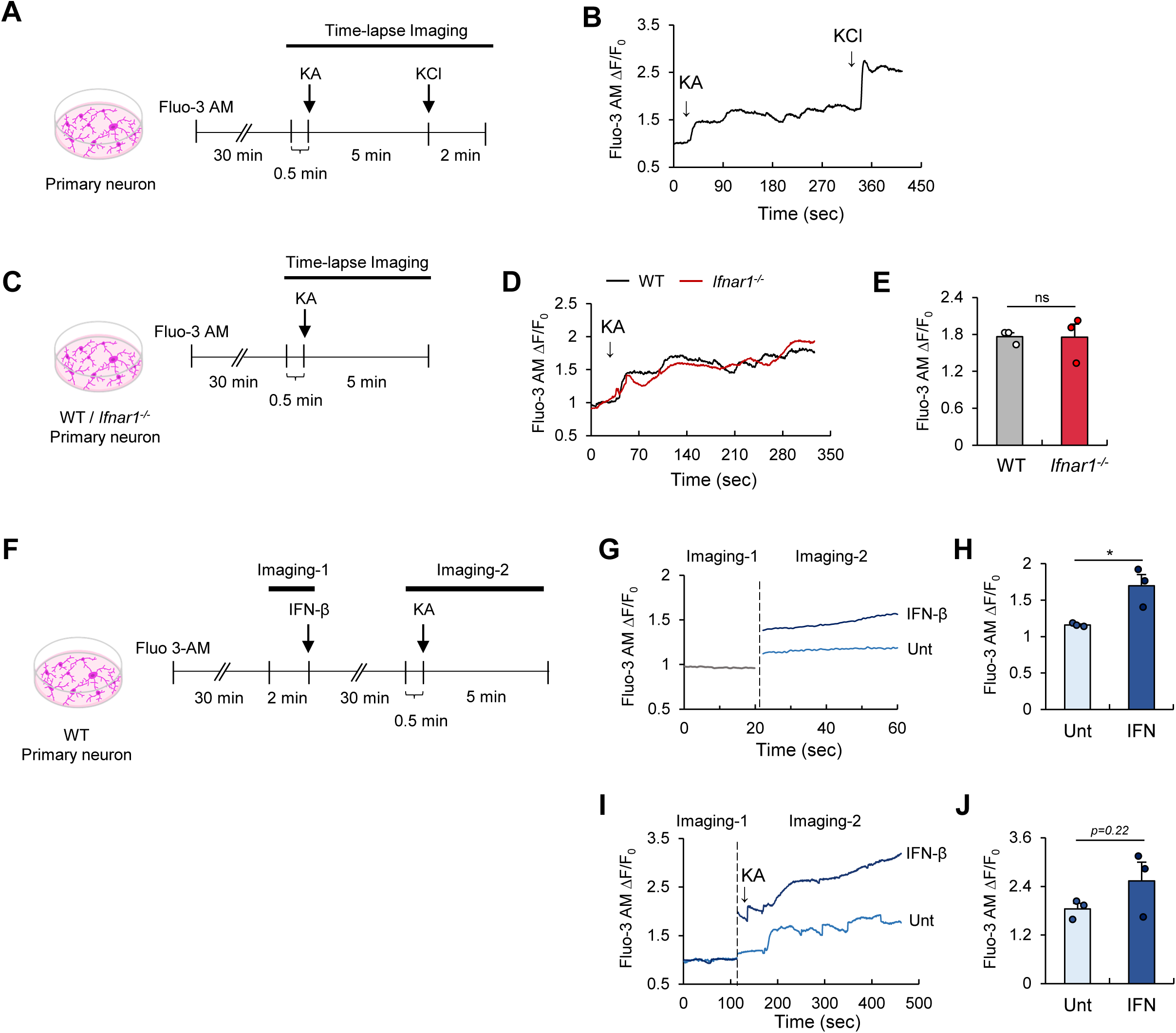
WT and *Ifnar1^-/-^* primary neurons do not show difference in excitability in response to kainic acid, while IFN-β priming significantly alters neuronal excitability. **(A, C, F)** Schematic overview of the experimental design. Primary neurons were loaded with the calcium indicator Fluo-3 AM and incubated for 30 min at 37 °C. Time-lapse imaging began with baseline activity recording, followed by sequential treatments with KA (1 mM) and KCl (50 mM). For IFN-β priming, IFN-β (5 ng/ml) was applied after the initial baseline measurement. Following a 30 min incubation with IFN-β, baseline alterations were assessed, and KA and KCl treatments were performed as previously described. **(B)** Representative calcium traces of primary neurons in responses to KA and KCl treatments. Only neurons responsive to KCl were included in the analysis. **(D)** Representative calcium traces of WT and *Ifnar1^-/-^* primary neurons, showing responses to KA treatment. **(E)** Maximum ΔF/F_0_ in response to KA treatment in WT and *Ifnar1^-/-^* primary neurons. **(G)** Representative calcium traces of WT primary neurons, with or without IFN-β priming. **(H)** Maximum ΔF/F_0_ in response to IFN-β treatment. **(I)** Representative calcium traces of WT primary neurons responding to KA, with or without prior IFN-β priming. **(J)** Maximum ΔF/F_0_ (F_0_, baseline before IFN-β treatment) in response to KA in untreated-versus IFN-β-primed neurons. *n* = 3 per group. Data are presented as mean ± SEM. Statistical analysis was performed using Student’s *t* test. * *P* < 0.05, ns; not significant.

We then examined whether IFN priming could alter baseline neuronal activity or affect the response to subsequent KA stimulation. After recording initial baseline activity, cells were incubated with IFN-β for 30 min, then imaging resumed to assess changes in baseline excitability and response to KA (Fig. 4F). Notably, IFN-β treatment significantly increased baseline neuronal excitability (Fig. 4G, H) and appeared to enhance the subsequent KA-induced response, though this enhancement did not reach statistical significance (Fig. 4I, J). These findings suggest that type I IFN signaling directly increases neuronal excitability.

### Type I interferon signaling modulates mTOR activation

The mammalian target of rapamycin (mTOR) pathway is a crucial signaling pathway that regulates cell growth, metabolism, proliferation, and survival, thereby maintaining cellular homeostasis^43^. Clinically, mTORopathies-a group of disorders caused by dysregulation of the mTOR pathway-are closely associated with epileptogenesis and drug-resistant epilepsy^43–47^. Furthermore, preclinical studies have shown that while many mTORopathy models develop epilepsy, mTOR activation is also observed in acquired epilepsy models^44,46–49^.

To determine whether the reduced seizure severity observed in *Ifnar1^-/-^* mice is accompanied by decreased mTOR activation, we first sought to establish the time points of mTOR activation following KA administration. In wild-type mice, we analyzed the hippocampus and cortex at various time intervals post-KA injection. Using the phosphorylation of S6 (a marker of mTORC1 activation) and phosphorylation of AKT (a marker of mTORC2 activation) as indicators, we found that both mTORC1 and mTORC2 were activated as early as 1 h after KA injection in the hippocampus. Additionally, mTORC1 activation in the cortex showed an increasing trend beginning 1 h post-injection (Fig. 5A, B). In this context, we compared mTOR activation between wild-type and *Ifnar1^-/-^* mice 1 h after KA administration (Fig. 5C). Notably, the activation of both mTORC1 and mTORC2 in the hippocampus was significantly attenuated in *Ifnar1^-/-^* mice compared to wild-type mice during seizures (Fig. 5D, E). A similar trend of reduced mTOR activation was observed in the cortex (Fig. 5D, F). These findings demonstrate that type I IFN signaling contributes to the activation of the mTOR pathway in the brain during seizures and suggest that the reduced seizure severity in *Ifnar1^-/-^* mice may be associated with attenuated hyperactivation of the mTOR pathway.

**Fig 5.**
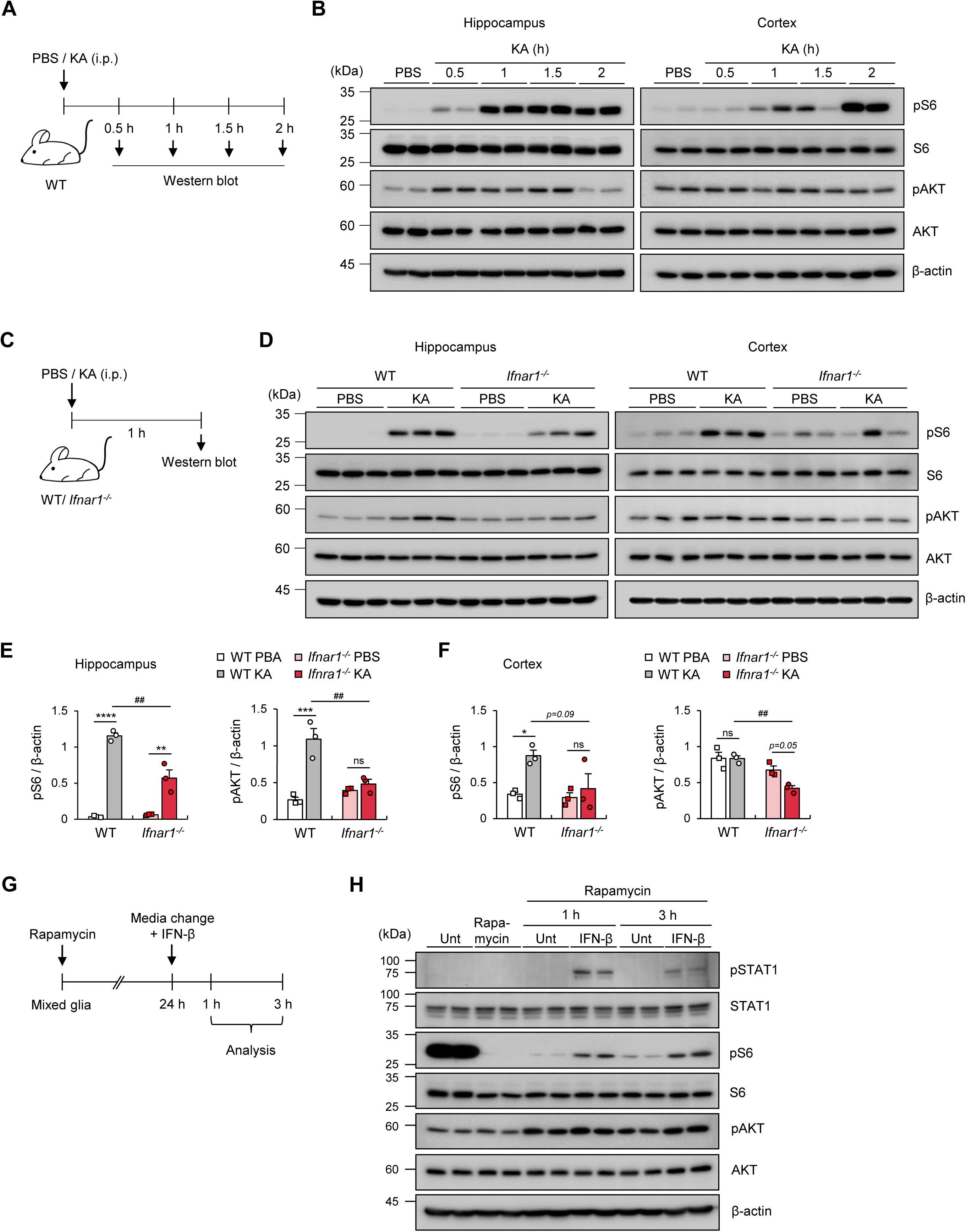
Attenuated seizure severity in *Ifnar1^-/-^* is associated with decreased mTOR pathway activation during kainic acid-induced seizures. **(A)** Schematic diagram illustrating KA-induced seizure in WT mice. Mice were harvested at various time points following KA injection, and the hippocampus and cortex were collected for western blot analysis. **(B)** Representative immunoblot images from the hippocampus and cortex of PBS- or KA- injected WT mice at various time points. **(C)** Schematic overview of KA-induced seizure in WT and *Ifnar1^-/-^* mice. **(D)** Representative western blot images from the hippocampus and cortex of PBS- or KA-injected WT and *Ifnar1^-/-^* mice 1 h after KA administration. **(E, F)** Quantification of pS6 and pAKT protein levels normalized to β-actin in the hippocampus and cortex of WT and *Ifnar1^-/-^* mice. *n* = 3 per group. **(G)** Schematic diagram of the mixed glia experiment. Mixed glia were treated with rapamycin (20 nM) for 24 h. The media were then replaced with fresh media, containing IFN-β (10 ng/ml) or without IFN-β. Cells were collected 1 h or 3 h later for western blot analysis. **(H)** Representative immunoblot images of mixed glia treated as in (G). Data are presented as mean ± SEM. Statistical analysis was performed using two-way ANOVA with Tukey’s post hoc test for (E, F). * *P* < 0.05, **,^##^ *P* < 0.01, *** *P* < 0.001, **** *P* < 0.0001, ns; not significant.

To further validate the impact of type I IFN signaling on mTOR activation, we conducted *in vitro* experiments using mixed glial cultures containing approximately 20% microglia and 5∼60% astrocytes (Fig. 5G and Supplementary Fig. 3). Since the mTOR pathway is typically activated under resting conditions, we first inhibited mTORC1 by incubating the cells with the mTORC1 inhibitor rapamycin for 24 h. After this treatment, we refreshed the media and applied IFN-β for either 1 or 3 h. Immunoblot analysis revealed that IFN-β treatment caused a robust S6 phosphorylation, an indication of mTORC1 activation, as well as STAT1 phosphorylation (Fig. 5H). These results demonstrate that type I IFN signaling activates mTOR pathway in the glial cells.

## Discussion

Type I interferon (IFN) has recently been recognized not only as an immune cytokine but also as an important neuromodulator within the central nervous system (CNS)^32,50^. Although its role in various CNS disorders is increasingly understood, the relationship between type I IFN signaling and seizure activity remains largely unexplored. In this study, we provide evidence that type I IFN signaling influences seizure activity, particularly affecting the initiation phase.

Our comparative analysis of seizure phenotypes between wild-type and *Ifnar1^-/-^* mice demonstrated that IFNAR1 deficiency reduced seizure severity in the acute phase, suggesting that type I IFN signaling may exacerbate seizure activity. In *Ifnar1^-/-^* mice, both neuronal activity and microglial activation were attenuated during the acute seizure phase compared to wild-type mice. Moreover, intracerebroventricular injection of IFN-β intensified seizure severity, indicating that the presence of type I IFN signaling worsens seizure conditions, while its absence (in *Ifnar1^-/-^* mice) provides a degree of protection. The role of type I IFN in modulating neuronal responses was further supported by experiments with primary neurons. While KA-induced responses did not significantly differ between wild-type and *Ifnar1^-/-^* neurons, IFN priming increased neuronal activity, suggesting that IFN-induced excitability, rather than inherent differences between wild-type and *Ifnar1^-/-^* neurons, may contribute to the observed seizure severity differences *in vivo*. However, the lack of glial interactions in this experiment may limit these findings. Given that neuron-glia communication plays a significant role in regulating neuronal excitability and seizure outcomes, this neuron-only comparison may not fully reflect the physiological dynamics.

Despite our findings, we cannot rule out the possibility that intrinsic developmental differences between wild-type and *Ifnar1^-/-^* mice contribute to the observed differences in seizure severity. Previous studies have demonstrated that the ablation of type I IFN signaling alters neuronal and glial properties^14,15,51^. For instance, aged male *Ifnar1^-/-^* mice exhibit elevated levels of *Gabrb1*, which encodes the GABA_A_ receptor, a critical component for inhibitory neurotransmission^51^. Additionally, type I IFN signaling influences microglial phagocytosis and modulates the expression of glutamate transporter in astrocytes^14,15^. These changes could impact neuronal and glial function, potentially contributing to the reduced seizure susceptibility observed in *Ifnar1^-/-^* mice. However, in our data (Fig. 4I, J), the lack of significant differences in KA-induced responses between wild-type and *Ifnar1^-/-^* neurons (Fig. 4I, J) suggests that the *in vivo* differences in seizure severity are more likely due to the presence of type I IFN signaling rather than intrinsic differences in neuronal excitability between the genotypes. Further investigation will be needed to clarify these points. Additionally, elucidating the specific cell types that contribute to the type I IFN-mediated augmentation of seizure progression will be an important next step.

Our study provides new insights into the role of type I IFN signaling in modulating the mTOR pathway, which is critical in regulating cellular growth, metabolism, neuronal excitability, and glia activation. Dysregulated mTOR signaling, particularly due to mutations in the *TSC1* or *TSC2*, is strongly linked to epilepsy. Previous research has shown that hyperactivation of mTOR specifically in microglia leads to spontaneous recurrent seizures, with elevated levels of IFN-α and IFN-β observed in both the hippocampus and cortex^52^. This evidence led us to hypothesize that the mTOR pathway may interact with type I IFN signaling to influence seizure severity. Our results demonstrated that mTOR activation during seizures was significantly reduced in *Ifnar1^-/-^* mice compared to wild-type mice, suggesting that type I IFN signaling may contribute to seizure pathogenesis by promoting mTOR activation, thereby potentially exacerbating neuronal hyperexcitability and microglia activation. This connection opens new avenues for exploring how immune signaling modulates key intracellular pathways in epilepsy.

In conclusion, our findings suggest that type I IFN signaling plays a multifaceted role in seizures, impacting both neuronal activity and microglial responses. The protective effects observed in *Ifnar1^-/-^* mice highlight the importance of further elucidating the mechanisms by which type I IFN exacerbates seizures, as these insights lead to novel therapeutic strategies for epilepsy. Given the limited research on this topic, future studies should aim to clarify the specific molecular and cellular interactions through which type I IFN’s contributes to seizure dynamics, particularly during the early stages of seizure initiation and propagation.

## Acknowledgements

This work was supported by the National Research Foundation of Korea Grant funded by the Korean Government (2022R1A2C2007569, RS-2023-00207834) and by a grant of the Korea Health Technology R&D Project through the Korea Health Industry Development Institute (KHIDI), funded by the Ministry of Health & Welfare, Republic of Korea (RS-2024-00405260).

## Author Contributions

J.-H.M. and J.-W.Y. conceived and designed the study. J.-C. E., C. L., I. H., and J.C. performed and analyzed the experiments. SSJ and CK provided critical materials and technical advice. J.-W.Y. supervised the entire project. J.-H.M. and J.-W.Y. wrote the manuscript. All authors read and approved the final version of the manuscript.

## Data availability

All data needed to evaluate the conclusions in the paper are present in the paper or the Supplementary Materials.

