## Supplementary Figures for "Type I interferon signaling enhances kainic acid-induced seizure severity"

### Supplementary Figure 1

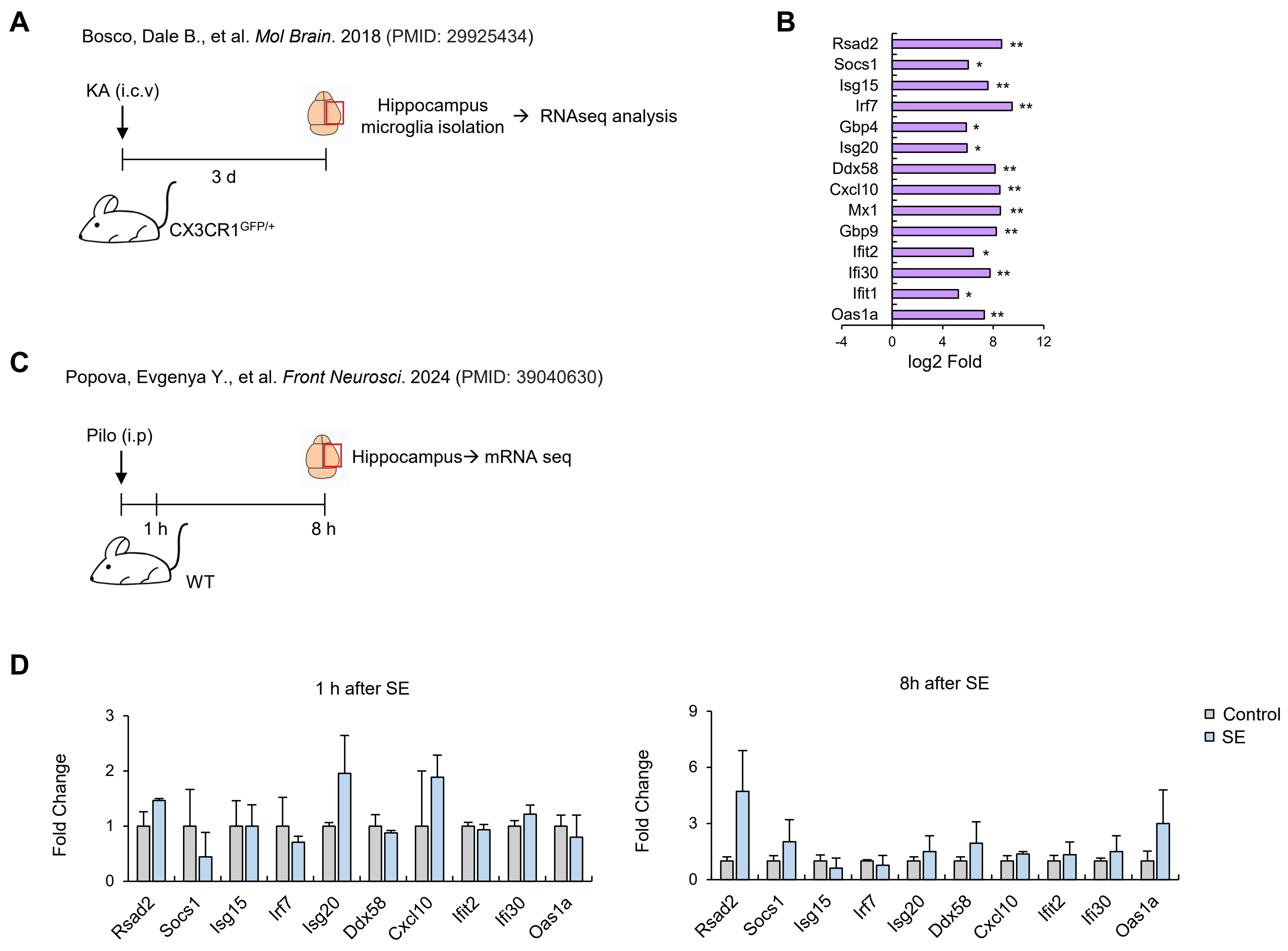


**Supplementary Figure 1. Upregulation of interferon-stimulated genes (ISGs) in a seizure mouse model. (A)** Re-analysis of previously published RNA-seq data to assess ISG expression changes after seizures. Microglia were isolated from the hippocampus 3 days post-KA injection and compared to sham controls. Excel file containing gene names, base mean expression (control and 3 days post-KA) with standard deviation, log2 fold change, and adjusted p-values was re-analyzed. ISGs were then selected, and their expression changes were evaluated. (**B**) Expression levels of ISGs in microglia from the hippocampus 3 days after KA administration compared to sham controls. RNA expression levels are presented as log2 fold changes relative to controls, with adjusted p-values. *n* = 3 per group.. **(C)** Re-analysis of additional RNA-seq data (GSE198498) to investigate ISG expression changes at earlier time points following seizures. Hippocampal tissue was harvested 1 h and 8 h post-status epilepticus (SE) induced by systemic pilocarpine injection and compared to controls. Commonly targeted ISGs were selected for analysis. (**D**) Expression levels of ISGs in the hippocampus 1 h and 8 h after SE compared to controls. RNA expression levels are shown as fold changes relative to controls. *n* = 2-3 per group.

#### Supplementary Figure 2

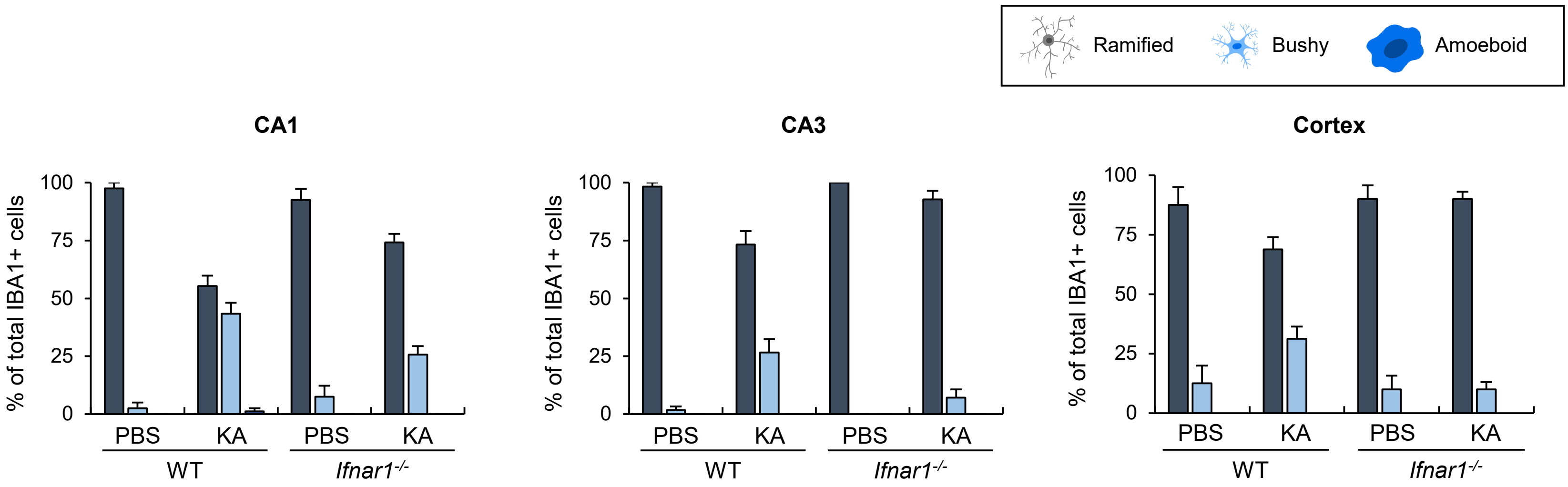


**Supplementary Figure 2. Differential microglial morphological changes in wild-type and *Ifnar1*$^{-/-}$ mice following kainic acid-induced seizures.** Microglia morphology was assessed in WT and *Ifnar1*$^{-/-}$ mice 2 hours after KA administration. Microglia were categorized into three subsets—ramified, bushy, and amoeboid—based on the ratio of the longest process length to soma diameter. The proportions of each morphological subset were compared between PBS- and KA-injected WT and *Ifnar1*$^{-/-}$ mice. PBS, $n$ = 5-6; KA, $n$ = 7-8.

**Supplementary Figure 3**

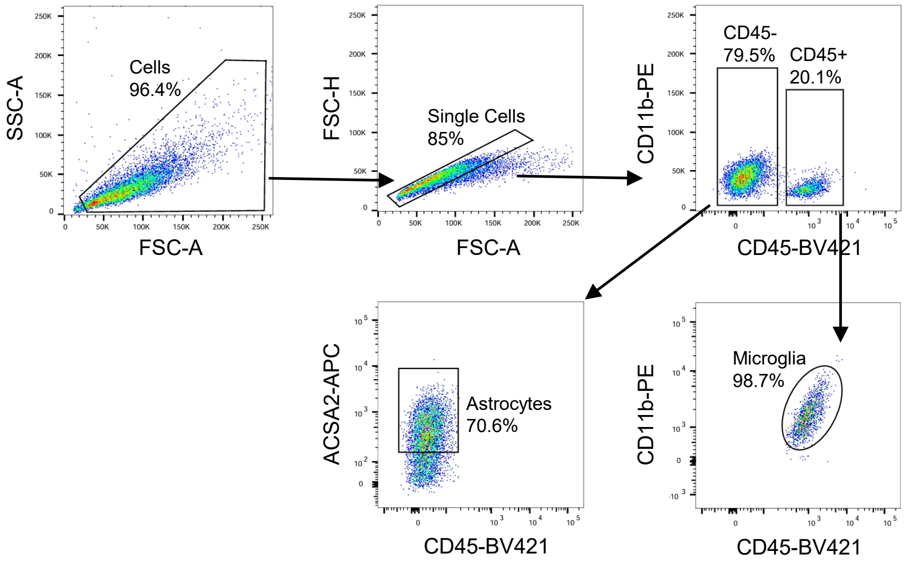


**Supplementary Figure 3. Composition of mixed glial cultures.** Mixed glial cultures were prepared from postnatal pups (P0–P2). The cultures consisted of approximately 20% microglia, 5–60% astrocytes, and other cell types. Microglia were identified as $CD45^{+}CD11b^{int}$ , and astrocytes were identified as $CD45^{-}ACSA2^{+}$ using flow cytometric analysis.

### Uncropped full immunoblots

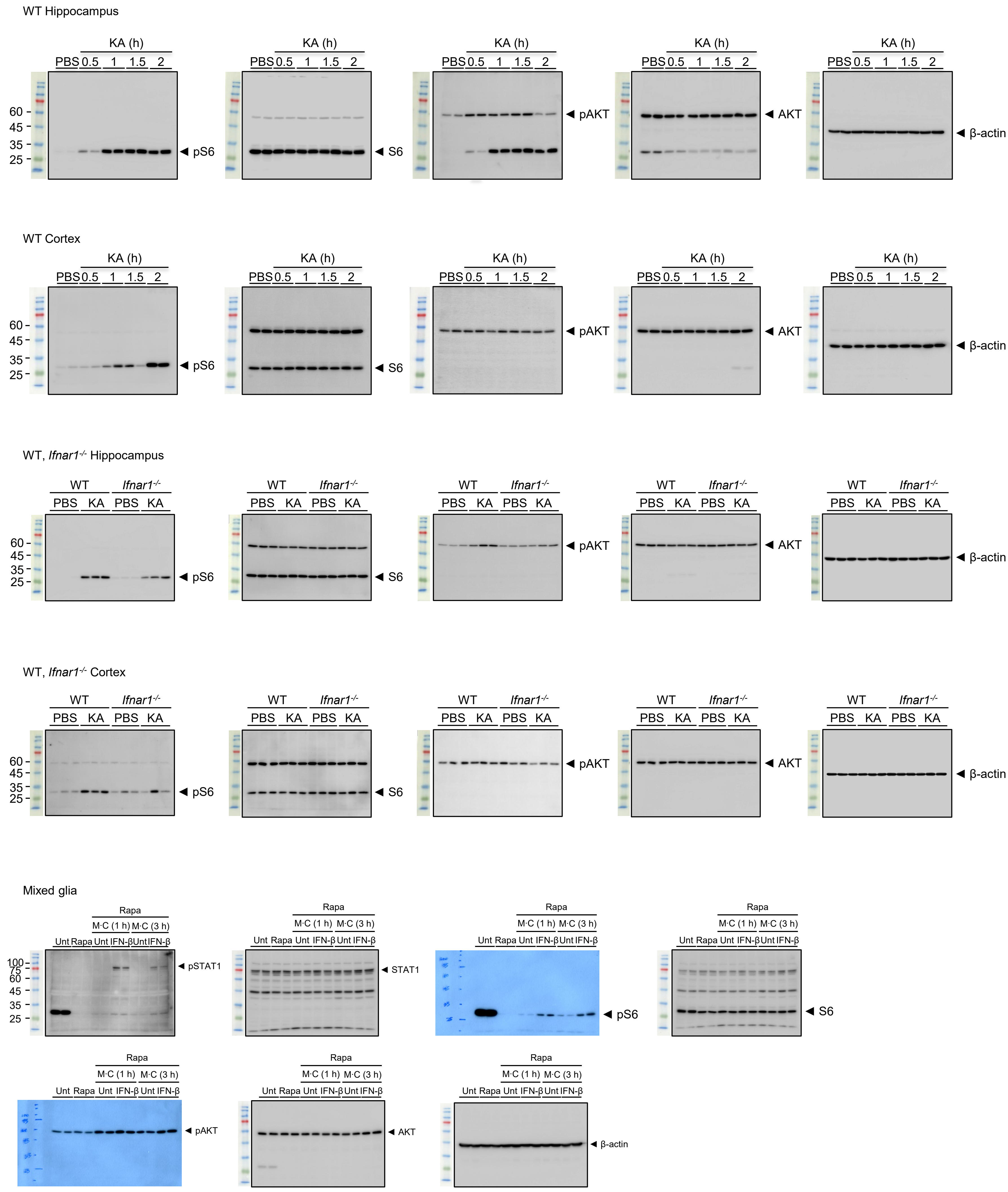
